# Microevolution of aquatic *Streptococcus agalactiae* ST-261 from Australia indicates dissemination via imported tilapia and ongoing adaptation to marine hosts or environment

**DOI:** 10.1101/301424

**Authors:** Minami Kawasaki, Jerome Delamare-Deboutteville, Rachel O Bowater, Mark J Walker, Scott Beatson, Nouri L Ben Zakour, Andrew C Barnes

**Affiliations:** The University of Queensland, School of Biological Sciences and Centre for Marine Science, St Lucia Campus, Brisbane, Queensland, Australia; Biosecurity Queensland (North Region), Department of Agriculture and Fisheries Townsville, Queensland, Australia; The University of Queensland, School of Chemistry and Molecular Biosciences and Australian Infectious Diseases Research Centre,, St Lucia Campus, Brisbane, Queensland, Australia

**Keywords:** Streptococcus agalactiae, evolution, genome analysis, epidemiology, fish, aquaculture

## Abstract

*Streptococcus agalactiae* (GBS) causes disease in a wide range of animals. The serotype 1b lineage is highly adapted to aquatic hosts, exhibiting substantial genome reduction compared with terrestrial con-specifics. Here we sequence genomes from 40 GBS isolates including 25 from wild fish and captive stingrays in Australia, six local veterinary or human clinical isolates, and nine isolates from farmed tilapia in Honduras and compare with 42 genomes from public databases. Phylogenetic analysis based on non-recombinant core genome SNPs indicated that aquatic serotype Ib isolates from Queensland were distantly related to local veterinary and human clinical isolates. In contrast, Australian aquatic isolates are most closely related to a tilapia isolate from Israel, differing by only 63 core-genome SNPs. A consensus minimum spanning tree based on core genome SNPs indicates dissemination of ST-261 from an ancestral tilapia strain, which is congruent with several introductions of tilapia into Australia from Israel during the 1970s and 1980s. Pan-genome analysis identified 1,440 genes as core with the majority being dispensable or strain-specific with non-protein-coding intergenic regions (IGRs) divided amongst core and strain-specific genes. Aquatic serotype Ib strains have lost many virulence factors during adaptation, but six adhesins were well conserved across the aquatic isolates and might be critical for virulence in fish and targets for vaccine development. The close relationship amongst recent ST-261 isolates from Ghana, USA and China with the Israeli tilapia isolate from 1988 implicates the global trade in tilapia seed for aquaculture in the widespread dissemination of serotype Ib fish-adapted GBS.

**Importance:** *Streptococcus agalactiae* (GBS) is a significant pathogen of humans and animals. Some lineages have become adapted to particular hosts and serotype Ib is highly specialized to fish. Here we show that this lineage is likely to have been distributed widely by the global trade in tilapia for aquaculture, with probable introduction into Australia in the 1970s and subsequent dissemination in wild fish populations. We report variability in the polysaccharide capsule amongst this lineage, but identify a cohort common surface proteins that may be a focus of future vaccine development to reduce the biosecurity risk in international fish trade.

## 1. Introduction

*Streptococcus agalactiae*, or Lancefield Group B *Streptococcus* (GBS), is a commensal and occasionally pathogenic bacterium with a very diverse host range. A common commensal in the urogenital tracts of humans, GBS is also a leading cause of morbidity in newborns causing meningitis, septicaemia and pneumonia (1–4). *S*. *agalactiae* can cause septicaemic infections in cattle, domestic dogs and cats, in camels, and reptiles and amphibians (5–8). In fish, disease outbreaks caused by *S*. *agalactiae* have substantial impact on the aquaculture industry, particularly the production of warm fresh water species such as tilapia (*Oreochromis* spp.)(9–12). Most outbreaks to date in freshwater farmed fish have resulted from infection by highly adapted strains of GBS with genomes that are 10-15% smaller than their terrestrial conspecifics (13). Unusually, *S*. *agalactiae* also causes significant mortality in wild aquatic animals including grouper, stingrays and mullet (5) suggesting further adaptation to marine as well as freshwater aquatic hosts.

Microevolution within a bacterial species can be driven by host or environmental adaptation (13, 14), permitting inference of the epidemiology of disease outbreaks and how pathogens may have transferred within and between geographic regions (14–16). This requires analysis of factors that evolve at sufficiently rapid pace to be informative over relatively short timespans. In GBS, capsular serotyping either with antibodies or by ‘molecular serotyping’ (sequencing of the capsular operon) has become a widely used method of typing for population studies (17–20) and, currently, *S*. *agalactiae* can be divided into ten capsular serotypes (Ia, Ib and II-IX) (18, 21). Determining capsular serotypes is also critical for vaccine formulation since CPS is highly immunogenic and can confer excellent protection against infections by the homologous CPS serotype (20, 22, 23). Further typing resolution is provided by multilocus sequence typing (MLST), a method that has been employed to great effect to conduct global population studies of isolates based on genetic variations amongst relatively slowly evolving housekeeping genes (17). Combining molecular serotyping and MLST in the analysis of *S*. *agalactiae* revealed that the majority of isolates associated with aquatic environments and hosts fall within serotypes Ia and Ib, in which Ia isolates belong to ST-7 in clonal complex (CC) 7 and ST-103 in CC103 (12, 18, 24–28). Serotype Ib strains isolated in Central and South America are ST-260 and ST-552 in CC552 (27, 29) and strains isolated in Australia, Israel, Belgium, China, Ghana, USA and Southeast Asia belong to ST-261 (5, 6, 9, 13, 29–31). Serotype III is commonly causative of disease in humans, but has also been isolated from fish in Thailand, China and recently in Brazil (26, 32–34).

Whilst capsular serotyping and MLST have been useful in inferring origin and dispersal of GBS subtypes, they do not display sufficient resolution to explore spread and evolution within individual sequence types nor can they reflect the complete genetic diversity of *S*. *agalactiae* (35). The rapid fall in cost of whole genome sequencing coupled with multiplexing and rapid development of open source bioinformatics tools has permitted much deeper analysis of evolution, host adaptation and epidemiological modelling within single bacterial species (36) including those from aquatic hosts (14, 15). Bacteria such as *S*. *agalactiae* that can colonise multiple host species often have greater genomic intraspecies diversity (37). In GBS, two major evolutionary trends have been implicated in rapid adaptation to new hosts, namely, acquisition of new genes by lateral gene transfer and genome reduction via gene loss integral to host specialisation (13, 38). For example, *S*. *agalactiae* Ia strains, GD201008-001 and ZQ0910 isolated from tilapia in China carry a 10 kb genomic island (GI), which is absent from their closely related human isolate A909. Moreover, this 10 kb GI bears many similarities with *Streptococcus anginosus* SK52/DSM 20563 genome sequence suggesting possible transfer from *S*. *anginosus* to GBS, with implications for virulence in Tilapia (13, 39). During fish host adaptation, serotype Ib strains have undergone reductive evolution resulting in 10-25% of their genomes being lost compared to terrestrial *S*. *agalactiae* isolates and serotype Ia piscine strains (13). Evolution of *S*. *agalactiae* by genome reduction is an ongoing process with a high number of pseudogenes present in GBS genomes from aquatic sources (13).

Evolution of *S*. *agalactiae* and adaptation to aquatic hosts is an incomplete and ongoing process, consequently sequencing the genomes from a few isolates is insufficient to understand the full potential genetic diversity of *S*. *agalactiae* as a species (35). The pan-genome or supra-genome of a bacterial species defines the full complement of genes, or the union of all the gene sets, within the species (35). This pan-genome is subdivided into its core genome, which includes all the genes that are present in all the strains of the same bacterial species and must therefore be responsible for essential biological functions to allow the species to survive, and the accessory genome containing species-specific genes that are unique to single strains or constrained to a cohort of strains within the species; these genes contribute to the diversity makeup of the species. The pan-genome of a species resolves the true genomic diversity of that species and permits the identification of gene cohorts that are essential to the species as a whole, along with gene complements in the accessory genome that permit host or habitat specialization (35). Moreover, by identifying potential antigens within the pan-genome that are conserved across all strains that infect a particular host type, vaccine targets can be specified that are likely to cross-protect (35, 40). Indeed, the first multicomponent protein-containing universal vaccine against human *S*. *agalactiae* was developed using a pan-genome reverse vaccinology approach by analysing eight human isolates to predict putative antigens that were conserved amongst those strains (41). Some antigens in this vaccine are in the accessory genome consequently it is important to analyse as large a dispensable genome as possible for vaccine development (35, 41).

The *S*. *agalactiae* pan-genome is now well-advanced but still “open” (new genes continue to be added with more sequenced genomes) and geographically constrained (35, 40). In the present study, we sequenced genomes of new aquatic *S*. *agalactiae* strains isolated from tilapia in Honduras and from wild and captive marine fish in Australia. We infer potential epidemiological distribution of ST-261 in aquatic hosts in Australia and show continuing adaptation to salt water fish. Moreover, we identify conserved surface proteins across the ST-260 and ST-261 sequence types that may have potential for incorporation into for aquaculture of important food fish species such as tilapia and grouper.

## 2. Materials and Methods

### 2.1. Bacterial strains and culture conditions

Forty *S*. *agalactiae* isolates comprising strains collected from fish in Honduras and Australia, along with reptiles, humans and other terrestrial mammals from Australia were chosen for sequencing (Table 1). Of these 40 isolates, 25 strains were collected from several species of fish in Queensland, Australia, two human clinical strains were from Queensland, Australia, one strain was isolated from saltwater crocodile (*Crocodylus porosus*) in the Northern Territory, Australia and three isolates were collected from domestic animals including cats, dogs and cattle in Australia. Additionally, nine isolates originating from disease in farmed tilapia in Honduras were sequenced during this study. All isolates were maintained at −80°C in Todd Hewitt broth (THB) (Oxoid) containing 25% glycerol as frozen stock. The isolates were recovered from stock on Columbia agar containing 5% sheep blood (Oxoid) for 24 h at 28°C. For liquid culture, the isolates were grown in THB for 18 h with low agitation at 28°C.

**Table 1:** *S*. *agalactiae* isolates and sequences used in this study

| Isolate | Origin | Year | Host | Serotype | ST | Size<br>(Mbp) | Accession no. |
| --- | --- | --- | --- | --- | --- | --- | --- |
| QMA0264 | QLD, AU | 2008 | *Epinephelus lanceolatus* | Ib | 261 | 1.8 | SAMN07998566 |
| QMA0266 | QLD, AU | 2008 | *Epinephelus lanceolatus* | Ib | 261 | 1.8 | SAMN07998567 |
| QMA0267 | QLD, AU | 2008 | *Epinephelus lanceolatus* | Ib | 261 | 1.8 | SAMN07998568 |
| QMA0268 | QLD, AU | 2009 | *Pomadasys kaaken* | Ib | 261 | 1.8 | SAMN07998569 |
| QMA0271 | QLD, AU | 2009 | *Arius thalassinus* | Ib | 261 | 1.8 | SAMN07998570 |
| QMA0273 | QLD, AU | 2009 | *Arius thalassinus* | Ib | 261 | 1.8 | SAMN07998571 |
| QMA0274 | QLD, AU | 2009 | *Liza vaigensis* | Ib | 261 | 1.8 | SAMN07998572 |
| QMA0275 | QLD, AU | 2009 | *Aptychotrema rostrata* | Ib | 261 | 1.8 | SAMN07998573 |
| QMA0276 | QLD, AU | 2009 | *Himantura granulata* | Ib | 261 | 1.8 | SAMN07998574 |
| QMA0277 | QLD, AU | 2009 | *Dasyatis fluviorum* | Ib | 261 | 1.8 | SAMN07998575 |
| QMA0280 | QLD, AU | 2010 | *Epinephelus lanceolatus* | Ib | 261 | 1.8 | SAMN07998576 |
| QMA0281 | QLD, AU | 2010 | *Epinephelus lanceolatus* | Ib | 261 | 1.8 | SAMN07998577 |
| QMA0284 | QLD, AU | 2010 | *Epinephelus lanceolatus* | Ib | 261 | 1.8 | SAMN07998578 |
| QMA0285 | QLD, AU | 2010 | *Epinephelus lanceolatus* | Ib | 261 | 1.8 | SAMN07998579 |
| QMA0287 | QLD, AU | 2010 | *Pomadasys kaaken* | Ib | 261 | 1.8 | SAMN07998580 |
| QMA0290 | QLD, AU | 2010 | *Arius thalassinus* | Ib | 261 | 1.8 | SAMN07998581 |
| QMA0292 | QLD, AU | 2010 | *Aptychotrema rostrata* | Ib | 261 | 1.8 | SAMN07998582 |
| QMA0294 | QLD, AU | 2010 | *Epinephelus lanceolatus* | Ib | 261 | 1.8 | SAMN07998583 |
| QMA0320 | QLD, AU | 2010 | *Dasyatis fluviorum* | Ib | 261 | 1.8 | SAMN07998584 |
| QMA0321 | QLD, AU | 2010 | *Dasyatis fluviorum* | Ib | 261 | 1.8 | SAMN07998585 |
| QMA0323 | QLD, AU | 2010 | *Dasyatis fluviorum* | Ib | 261 | 1.8 | SAMN07998586 |
| QMA0326 | QLD, AU | 2010 | *Dasyatis fluviorum* | Ib | 261 | 1.8 | SAMN07998587 |
| QMA0347 | QLD, AU | 2010 | *Dasyatis fluviorum* | Ib | 261 | 1.8 | SAMN07998588 |
| QMA0368 | QLD, AU | 2010 | *Epinephelus lanceolatus* | Ib | 261 | 1.8 | SAMN07998589 |
| QMA0369 | QLD, AU | 2011 | *Epinephelus lanceolatus* | Ib | 261 | 1.8 | SAMN07998590 |
| QMA0485 | Honduras | 2014 | *Oreochromis niloticus* | Ib | 260 | 1.8 | SAMN07998597 |
| QMA0487 | Honduras | 2014 | *Oreochromis niloticus* | Ib | 260 | 1.8 | SAMN07998598 |
| QMA0488 | Honduras | 2014 | *Oreochromis niloticus* | Ib | 260 | 1.8 | SAMN07998599 |
| QMA0489 | Honduras | 2014 | *Oreochromis niloticus* | Ib | 260 | 1.8 | SAMN07998600 |
| QMA0494 | Honduras | 2014 | *Oreochromis niloticus* | Ib | 260 | 1.8 | SAMN07998601 |
| QMA0495 | Honduras | 2014 | *Oreochromis niloticus* | Ib | 260 | 1.8 | SAMN07998602 |
| QMA0496 | Honduras | 2014 | *Oreochromis niloticus* | Ib | 260 | 1.8 | SAMN07998603 |
| QMA0497 | Honduras | 2014 | *Oreochromis niloticus* | Ib | 260 | 1.8 | SAMN07998604 |
| QMA0499 | Honduras | 2014 | *Oreochromis niloticus* | Ib | 260 | 1.8 | SAMN07998605 |
| QMA0355 | QLD, AU | 2011 | *Homo sapiens* | Ia | 23 | 2.0 | SAMN07998591 |
| QMA0357 | QLD, AU | 2011 | *Homo sapiens* | Ia | 23 | 2.0 | SAMN07998592 |
| QMA0336 | NT, AU | 2005 | *Crocodylus porosus* | Ia | 23 | 2.0 | SAMN07998593 |
| QMA0300 | QLD, AU | 2008 | *Canis lapis familiaris* | V | 1 | 2.1 | SAMN07998594 |
| QMA0303 | QLD, AU | 2009 | *Felis catus* | V | 1 | 2.1 | SAMN07998595 |
| QMA0306 | QLD, AU | 2005 | *Bos taurus* | V | 1 | 2.2 | SAMN07998596 |
| GS16-0008 | Ghana | 2016 | *Oreochromis niloticus* | Ib | 261 | 1.8 | SRX2698682 |
| GS16-0031 | Ghana | 2016 | *Oreochromis niloticus* | Ib | 261 | 1.8 | SRX2698681 |
| GS16-0035 | Ghana | 2016 | *Oreochromis niloticus* | Ib | 261 | 1.8 | SRX2698680 |
| GS16-0046 | Ghana | 2016 | *Oreochromis niloticus* | Ib | 261 | 1.8 | SRX2698679 |
| ND2-22 | Israel | 1988 | *Oreochromis niloticus* | Ib | 261 | 1.8 | FO393392 |
| 138P | USA | 2007 | *Oreochromis niloticus* | Ib | 261 | 1.8 | CP007482 |
| 138spar | USA | 2011 | *Oreochromis niloticus* | Ib | 261 | 1.8 | CP007565.1 |
| GX026 | China | 2011 | *Oreochromis niloticus* | Ib | 261 | 1.8 | CP011328.1 |
| S13 | Brazil | 2015 | *Oreochromis niloticus* | Ib | 552 | 1.8 | CP018623.1 |
| S25 | Brazil | 2015 | *Oreochromis niloticus* | Ib | 552 | 1.8 | CP015976.1 |
| SA20 | Brazil |  | *Oreochromis niloticus* | Ib | 552*** | 1.8 | CP003919.2 |
| GD201008-001 | China | 2010 | *Oreochromis niloticus* | Ia | 7 | 2.1 | NC_018646.1 |
| HN016 | China | 2011 | *Oreochromis niloticus* | Ia | 7 | 2.1 | NZ_CP011325.1 |
| WC1535 | China | 2015 | *Oreochromis niloticus* | Ia | 7 | 2.2 | NZ_CP016501.1 |
| A909 | USA |  | *Homo sapiens* | Ia | 7 | 2.1 | NC_007432.1 |
| GBS85147 | Brazil |  | *Homo sapiens* | Ia | 103 | 2.0 | NZ_CP010319.1 |
| Sag37 | China | 2014 | *Homo sapiens* | Ib* | 12 | 2.2 | NZ_CP019978.1 |
| GBS1-NY | USA | 2012 | *Homo sapiens* | II | 22 | 2.2 | NZ_CP007570.1 |
| GBS2_NM | USA | 2012 | *Homo sapiens* | II | 22 | 2.2 | NZ_CP007571.1 |
| GBS6 | USA | 2009 | *Homo sapiens* | II | 22 | 2.2 | NZ_CP007572.1 |
| FDAARGOS_254 | USA | 2014 | *Homo sapiens* | II* | 22 | 2.2 | CP020449.1 |
| COH1 | USA |  | *Homo sapiens* | III | 17 | 2.1 | NZ_HG939456.1 |
| NEM316 | France |  | *Homo sapiens* | III | 23 | 2.2 | NC_004368.1 |
| CU_GBS_08 | Hong Kong | 2008 | *Homo sapiens* | III | 283 | 2.1 | NZ_CP010874.1 |
| CU_GBS_98 | Hong Kong | 1998 | *Homo sapiens* | III | 283 | 2.0 | NZ_CP010875.1 |
| NGBS128 | Canada | 2010 | *Homo sapiens* | III | 17 | 2.1 | NZ_CP012480.1 |
| SG-M1 | Singapore | 2015 | *Homo sapiens* | III | 283 | 2.1 | NZ_CP012419.2 |
| H002 | China | 2011 | *Homo sapiens* | III | 736 | 2.1 | NZ_CP011329.1 |
| Sag158 | China | 2014 | *Homo sapiens* | III* | 19 | 2.1 | NZ_CP019979.1 |
| NGBS061 | Canada | 2010 | *Homo sapiens* | IV | 459 | 2.2 | NZ_CP007631.1 |
| NGBS572 | Canada | 2012 | *Homo sapiens* | IV | 452 | 2.1 | NZ_CP007632.1 |
| 2603V/R | USA |  | *Homo sapiens* | V | 110 | 2.2 | NC_004116.1 |
| CNCTC10/84 | USA |  | *Homo sapiens* | V | 26 | 2.0 | NZ_CP006910.1 |
| NGBS357 | Canada | 2011 | *Homo sapiens* | V | 1 | 2.2 | NZ_CP012503.1 |
| SS1 | USA | 1992 | *Homo sapiens* | V | 1 | 2.1 | NZ_CP010867.1 |
| GBS-M002 | China | 2014 | *Homo sapiens* | VI* | 1 | 2.1 | NZ_CP013908.1 |
| SA111 | Portugal | 2013 | *Homo sapiens* | II* | 61** | 2.3 | NZ_LT545678.1 |
| FWL1402 | China | 2014 | *Hoplobatrachus chinensis* | III* | 739** | 2.1 | NZ_CP016391.1 |
| 09mas018883 | Sweden |  | *Bos taurus* | V* | 1 | 2.1 | NC_021485.1 |
| GBS ST-1 | USA | 2015 | *Canis lapis familiaris* | V | 1 | 2.2 | NZ_CP013202.1 |
| ILRI005 | Kenya |  | *Camelus dromedarius* | V* | 609 | 2.1 | NC_021486.1 |
| ILRI112 | Kenya |  | *Camelus dromedarius* | VI* | 617 | 2.0 | HF952106.1 |

AU, Australia; QLD, Queensland; NT, Northern Territory; ST, Sequence Type.
\*Serotype was detected with Kaptive. \*\* ST was determined via Center for Genomic Epidemiology.
\*\*\* gap in *glcK*

### 2.2. DNA extraction and sequencing

Genomic DNA (gDNA) was extracted from 10 ml early-stationary phase THB cultures with the Qiagen DNeasy mini kit (Qiagen) according to the manufacturer’s instructions. The quantity of extracted DNA was measured by Qubit fluorimetry (Invitrogen) and the quality was checked by agarose gel electrophoresis. To confirm the purity of the gDNA, the 16S rRNA gene was amplified by polymerase chain reaction (PCR) using universal primers 27F and 1492R (42) and the PCR products were sent to Australian Genome Research Facility (AGRF, Brisbane) for Sanger sequencing. The 16S amplicon sequences were assembled in Sequencher V5.2.2 and analysed by BLAST. Once identity and purity were confirmed, Nextera XT paired-end libraries were generated using gDNA from each isolate and sequenced on the Illumina HiSeq2000 platform system (AGRF, Melbourne).

### 2.3. De novo *assembly and annotation*

Illumina sequencing yielded between 5,288,952 and 12,577,340 read pairs for each strain. Read quality control and contaminant screening were performed using FastQC (43). Reads were trimmed using the clip function in Nesoni (https://github.com/Victorian-Bioinformatics-Consortium/nesoni) then assembled *de novo* with SPAdes Assembler version 3.11 (44). The assemblies of fish isolates in Queensland comprised about 1.8 Mbp of assembled sequence while terrestrial strains comprised 2 Mbp. The assembled contigs for all Queensland strains were reordered against an internal curated reference genome from *S*. *agalactiae* strain QMA0271, using Mauve contig ordering tool (45). Automated annotation was performed using Prokka 1.12 (46).

### 2.4. Molecular serotyping and multilocus sequence typing (MLST)

Reference sequences for the nine CPS serotypes (Table 2) (21) were retrieved from GenBank to generate a database for prediction of capsular serotype from the draft genomes with Kaptive using default settings (47). To determine multilocus sequence types (MLST), all draft assemblies were analysed using the Center for Genomic Epidemiology web-tools MLST ver. 1.8, using *S*. *agalactiae* configuration (available at https://cge.cbs.dtu.dk7/services/MLST/) (48).

**Table 2:** Reference sequences used for molecular capsule serotyping.

| Serotype | Accession Number | size (bp) | Reference |
| --- | --- | --- | --- |
| Ia | AB028896.2 | 25021 | (83) |
| Ib | AB050723.1 | 9987 | (84) |
| II | EF990365.1 | 12864 | (85) |
| III | AF163833.1 | 17276 | (86) |
| IV | AF355776.1 | 17596 | (87) |
| V | AF349539.1 | 18239 | (87) |
| VI | AF337958.1 | 16448 | (87) |
| VII | AY376403.1 | 14202 | (88) |
| VIII | AY375363.1 | 12637 | (88) |

### 2.5. Phylogenetic analysis

To estimate approximate phylogenetic relationship among our strains and other isolates with whole genome sequence available in GenBank (Table 1), a core genome single nucleotide polymorphism (SNP)-based phylogenetic tree was constructed. Whole genome sequences of Queensland and Honduras tilapia isolates, terrestrial isolates and the genomes obtained from GenBank were aligned with Parsnp in the Harvest Tools suite version 1.2 (49). The genome of *Streptococcus pyogenes* M1 GAS (Accession number: NC_002737.2) was also included as an outgroup for tree rooting. Hypothetical recombination sites in the core genome alignment were detected and filtered out with Gubbins (50). Maximum likelihood phylogenetic trees were inferred with RAxML version 8.2.9 (51) based on non-recombinant core genome SNPs under general time-reversible nucleotides substitution model (GTR) with 1000 bootstrap replicates. Ascertainment bias associated with analysis of only variable sites was accounted for using Felsenstein’s correction implemented in RAxML (51). The resulting tree was exported, rooted and nodes with low bootstrap supported collapsed with Dendroscope, and the resulting tree/cladogram annotated with Evolview V2 (52).

In order to infer possible phylogenetic relationships within the aquatic ST-261 clade of GBS, minimum spanning trees were constructed in MSTGold 2.4 (53) from a distance matrix based on core genome SNPs derived by alignment of 27 ST-261 genomes in Geneious V9.1 (Biomatters Inc). A consensus tree was constructed based on inference of 2400 trees and only those edges occurring in greater than 50% of trees were included in the consensus.

### 2.6. Pan-genome analysis

A reference pan-genome was built with GView server (54) using our curated genome of Queensland ST-261 serotype Ib grouper isolate, QMA0271 as a seed and 38 complete genomes from public databases added sequentially (Table 1). Only complete genomes were included into the reference pan-genome to avoid incomplete genes associated with the high fragmentation of draft sequences. For visualisation, draft and complete genomes were compared with the reference pangenome using BRIG 0.95 (55). To investigate core and accessory protein-coding genes, sequences from all strains were analysed with Roary (56) using default settings. Since Roary only considers protein-coding sequences, we used Piggy (57) to examine non protein-coding intergenic regions (IGRs), which comprise about 15% of the GBS genomes.

### 2.7. Identification and comparison of virulence factors

Virulence factor screening was performed using SeqFindr (58) by comparing the assembled genomes of all strains in this study along with the genomes available through GenBank (Table 1) against a list of 51 *S*. *agalactiae*-specific virulence factors sequences collected from the Virulence Factors Database (VFDB) (59), complemented by six additional sequences identified in the sequenced ST-261 *S*. *agalactiae* strain ND2-22 (Table 1).

### 2.8. Analysis of effects of SNPs in ST-261 clade

Putative effects of SNPs amongst and between the ST-261 clade were determined using SNPeff version 4.3p (60). First, a new *S*. *agalactiae* database was constructed from the Genbank-formatted curated reference genome for QMA0271 in accordance with the manual (http://snpeff.sourceforge.net/SnpEff_manual.html#databases). Then a VCF file, generated from curated SNPs generated by alignment of complete genomes of 27 ST-261 isolates in Geneious version 9.1, was annotated for SNP effect with SNPeff using the database from QMA0271 as reference (Supplementary information).

## 3. Results and Discussion

### 3.1. GBS isolates from marine fish and rays in Queensland and tilapia in Honduras belong to differing host-adapted lineages

The average size of draft genomes of isolates from fish and rays in Queensland was 1,801,022 bp, containing 1,881 genes on an average, while the mean genome size of strains from Honduran tilapia was 1,801,133 bp with an average gene number of 1,869, consistent with the small genomes associated with the host-adapted aquatic lineage (13). Molecular serotyping indicated that all Queensland and Honduras aquatic strains belong to serotype Ib but Queensland isolates belong to sequence type (ST)-261 while Honduras strains are ST-260. Both ST-260 and ST-261 have been identified infecting aquatic animals previously with ST-260 isolates found in tilapia from Brazil and Costa Rica and occupying clonal complex (CC) 552 along with ST-552 and ST-553 strains also isolated from tilapia in Latin America (27), whilst ST-261 has been isolated from tilapia in USA, China, Ghana and Israel (9, 13, 30).

The current isolates represent the first aquatic isolates sequenced from marine fish and stingrays indicating a wider host range (rays and marine finfish), environmental (freshwater to marine) and geographic distribution of ST-261 than previously observed (5, 6, 61). GBS strains isolated from humans and terrestrial animals in Queensland and Northern Territory, Australia were also sequenced and have larger genomes of 2,072,596 bp comprising 2,067 genes on average, suggesting that recent possible local transfer from terrestrial origin to Australian fish is highly unlikely, although probable transfer between humans and fish has been reported for ST-7 GBS elsewhere (8, 39, 62). Non-human mammalian strains from Australia sequenced in this study QMA0300 and QMA0303 belong to ST-1 serotype V, and QMA0306 belong to ST-67 serotype III, whereas human isolates, QMA0355 and QMA0357, and crocodile strain QMA0336 belong to ST-23 serotype Ia. Indeed, the high sequence identity between the human ST-23 serotype Ia isolates and those from farm-raised crocodiles are supportive of probable human transfer to these animals that has been implied previously (63).

To better inform possible origin of the ST-261 isolates from marine fish in Australia, a phylogenetic tree was constructed by maximum likelihood from 29,689 non-recombinant core genome SNPs and short indel positions derived by alignment of whole genome sequences of 25 Queensland fish isolates, 9 Honduras isolates, 6 Queensland terrestrial isolates and 42 genomes from public databases (Fig. 1A). Two distinct groups were resolved, the first comprising entirely of aquatic isolates (including serotype Ib isolates from ST552, ST-260 and ST-261) while the second major group comprised various isolates of terrestrial origin and some fish isolates from ST7 serotype Ia that may have infected fish via transfer from terrestrial sources (Fig. 1A). The serotype Ib aquatic group branched into three distinct lineages based on ST (Fig. 1A). One lineage comprised all Honduras strains of ST-260, which were derived from a lineage comprising ST-552 isolates from Brazil (Fig. 1A). This is supportive of previous observations in which an extended typing system based on MLST, virulence genes and serotype indicated geographic endemism within fish isolates from differing regions of Brazil (64). The isolates belonging to ST-261 from USA, Israel, Ghana, China and Queensland clustered together (Fig. 1A). The second major division, containing serotype Ia fish strains along with terrestrial isolates, was more complex, but isolates largely clustered in line with serotype and ST (Fig. 1A). The aquatic serotype Ia isolates clustered with human isolates of ST-7 (Fig 1A). These fish isolates have acquired a 10kb genomic island, putatively from *S*. *anginosus*, in contrast to their human ST7 serotype Ia relatives (39). Lineage 10 comprised serotype III ST-17 human isolates (Fig. 1A). Serotype Ia strains were divided amongst several further ST groups: Three Australian serotype Ia ST-23 strains (QMA0336, QMA0355 and QMA0357) isolated from humans and a saltwater crocodile clustered together with strains of serotype III ST-23 (NEM316) and serotype IV ST-452 (NGBS572) also of human origin (Fig. 1A). This lineage appears to be derived from NEM316 which is a frequent cause of late-onset disease in human infants (65). A further serotype Ia isolate was located in ST-103 and clustered together with two strains of serotype V ST-609 and ST-617 isolated from camels (66) (Fig. 1A). Two newly sequenced Australian terrestrial strains, QMA0300 isolated from a dog and QMA0303 isolated from a cat clustered with other serotype V ST-1 strains isolated from humans, cattle and dog hosts (Fig. 1A). This lineage also contained NGBS061 serotype IV ST-495 and GBS-M002 serotype VI ST-1 (Fig. 1A). ST-1 emerged as a significant cause of infection and disease in humans during the 1990s, but was recently inferred to have evolved from strains causing mastitis in cattle in the 1970s (67). Moreover, QMA0306 ST-1 serotype V from cattle in Queensland was closely related to SA111 ST-61 serotype II, which represents a host-adapted lineage of *S*. *agalactiae* that is dominant in cattle in Europe (68)(Fig. 1A).

**Figure 1.**
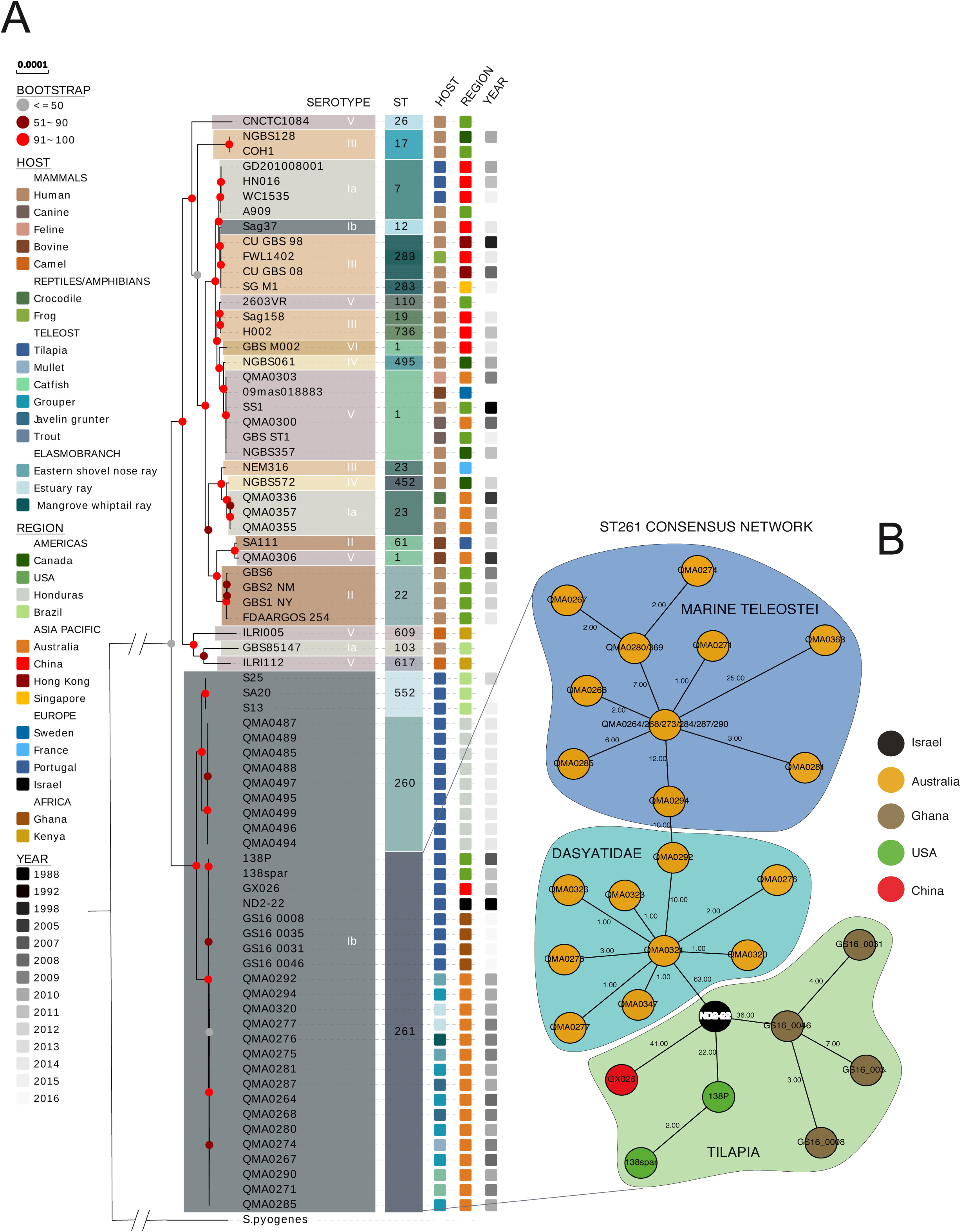
**A)** Maximum likelihood phylogeny of 82 *S*. *agalactiae* strains. The tree was inferred from alignment of 6050 non-recombinant core genome SNPs. Branch length was adjusted for ascertainment bias using Felsenstein’s correction implemented in RAxML (51). Nodes are supported by 1000 bootstrap replicates. The tree was rooted using *S*. *pyogenes* M1 GAS (Accession number: NC_002737.2) as an outgroup. B) Minimum spanning tree showing relationship amongst ST-261 serotype lb GBS isolates based on a distance matrix derived from non-recombinant core-genome SNP alignment. The consensus network was computed in MST-Gold (53).

Our phylogeny based on whole genome SNPs does not support recent direct transfer of GBS from Australian human clinical or terrestrial animal sources to marine fish and stingrays, in spite of close proximity of many of the wild fish cases to human habitation (6). Consequently, we refined our analyses to the ST-261 aquatic host-adapted lineage to attempt to infer possible route of introduction and subsequent evolution in Australian marine fish. A consensus minimum spanning tree based on a distance matrix comprised of all core genome SNPs derived from alignment of the ST-261 serotype Ib strains revealed a likely original introduction via tilapia from Israel, with only 63 core genome SNPs separating an Australian stingray isolate from the type strain of *S*. *difficile* (re-assigned as *S*. *agalactiae* serotype Ib (69)), isolated from tilapia in Israel in 1988 (Fig 1B)(70). Tilapia were imported on a number of occasions during the 1970s and 1980s from Israel into North Queensland around Cairns and Townsville, and a number of strains and hybrids have since colonised rivers and creeks throughout Queensland (71). Globally, the aquatic ST-261 lineage appears to have been transferred through human movements of tilapia for aquaculture and other purposes over the last several decades. The US and Chinese tilapia isolates also appear to derive from the early ND2-22 isolate, as do recent isolates from tilapia in Ghana (Fig 1B). Indeed, phylogenetic analysis by maximum likelihood of draft whole genome alignments suggest that the Ghanaian and Chinese isolates share a recent common ancestor that is derived from ND2-22, with only 60 SNPs separating ND2-22 and the Ghanaian strains (72). We identified only 36 core genome SNPs separating the Ghanaian isolates from ND2-22 but this reflects the smaller core genome in our study as a result of the high number of GBS isolates analysed (40 in the present study compared with 9 in the previous study (72).

The minimum spanning tree implicates continued adaptation of the ST-261 lineage post introduction and suggests that grouper (marine Teleostei family) may have been infected via estuary stingrays (Dasyatidae family) (Fig 1B). Stingrays are occasional prey for adult giant Queensland grouper and stingray barbs have been found in the gut of grouper post-mortem (Bowater, unpublished observation). ST-261 GBS has also caused mortality in captive stingrays in South East Queensland, translocated from Cairns in North Queensland (61)

### 3.2. The S. agalactiae pan-genome comprises a small core of protein-coding genes and is open

To further elucidate adaptation amongst the fish pathogenic GBS types a pan-genome was built from 39 complete genomes retrieved from GenBank and using our curated ST-261 grouper isolate QMA0271 as a high-quality reference seed genome. The resulting pan-genome was 4,074,275 bp (Fig. 2A). All-vs-all BLAST analysis of the genomes implemented in BRIG clearly indicated the substantial reduction in genome size amongst the fish pathogenic ST-261 cohort, as previously reported for a limited number of ST-261 isolates (13). Here, we find that ST-260 and ST552 fish-pathogenic sequence types within serotype Ib are similarly reduced and that conservation amongst the serotype Ib strains is high (Figure 2A). In total 4,603 protein-coding genes were predicted in *S*. *agalactiae* pan-genome using Roary (Fig. 2B), which is consistent with previous research (39). The number of core genes was 1,440 (representing 35% of the pan-genome) while previous studies reported 1,202 to 1,267 genes in the pan-genome (39, 40). These differences may result from the methods being used to examine the pan-genome, the difference in sequences number being used to create the pan-genome and finally the use of draft sequences in the analysis (39). A majority of protein-coding genes found in the pan-genome belonged to both the dispensable and strain-specific genes. This could be a result of the inclusion of a high number of serotype Ib strain sequences, which were all significantly smaller in size (approx. 1.8 Mbp) when compared with other isolates. Liu et al. (2013) demonstrated that removing Ib piscine isolates from their analysis resulted in an increase in the number of core genes.

**Figure 2.**
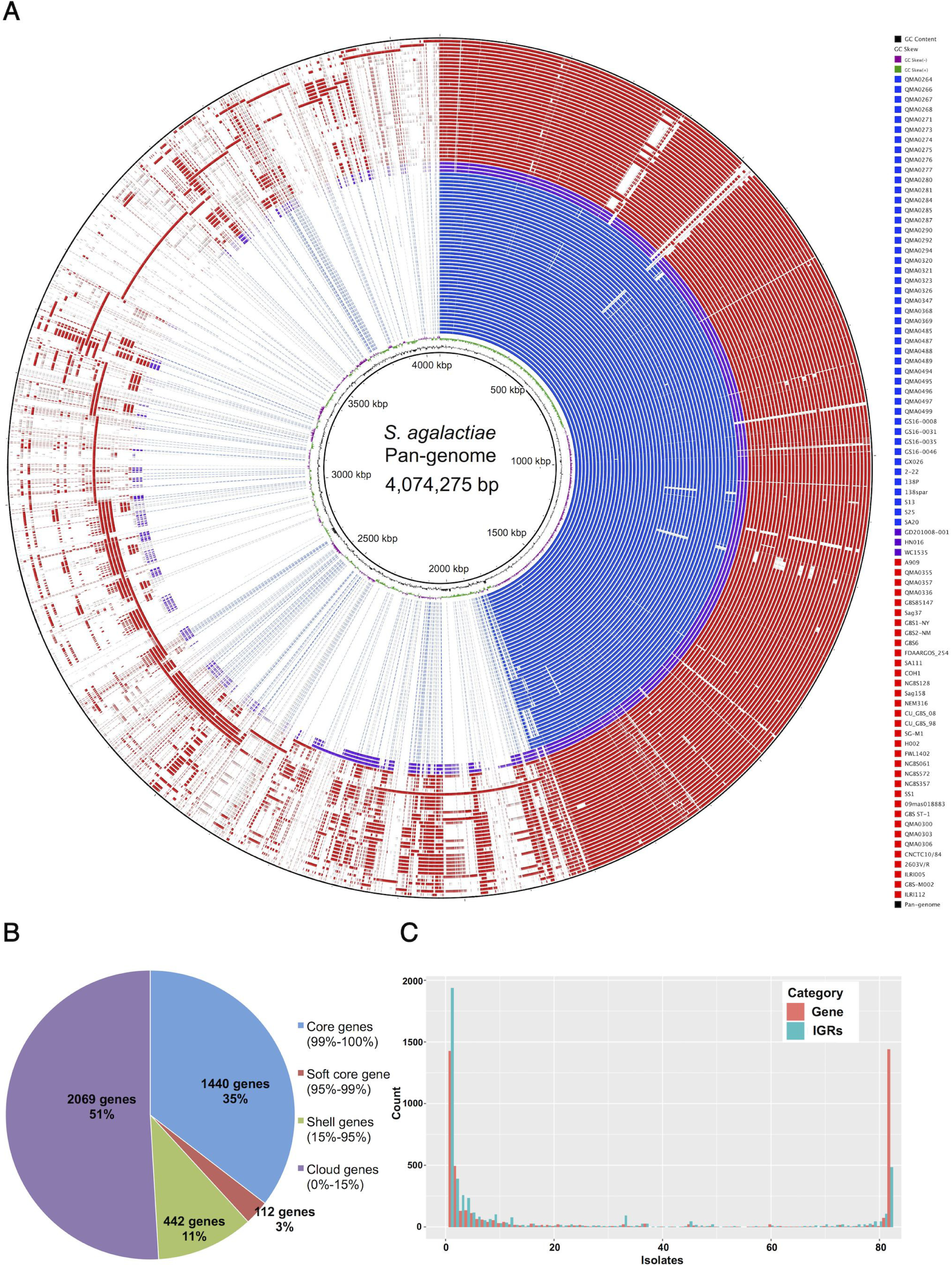
The pan-genome of *S*. *agalactiae*. A) BLASTN-based sequence comparison of 82 *S*. *agalactiae* genomes against the *S*. *agalactiae* pan-genome as reference constructed with BRIG 0.95 (55). Rings from the innermost to the outermost shows GC content and GC skew of the pan-genome reference, then sequence similarity of each of the 82 strains listed in the legend, from top to bottom rings are coloured according to origin with fish isolates belonging to ST-261, ST-260 and ST-552 serotype Ib in blue, fish strains belonging to serotype Ia in purple and terrestrial strains in red. The outermost ring (black) represents reference pan-genome. B) Proportion of protein-coding genes in the core, soft core, shell and cloud of the pan-genome of 82 *S*. *agalactiae* isolates determined with Roary. C) Histogram indicates the frequency of genes (protein-coding) in red and IGRs (non-protein-coding intergenic regions) in blue-green from 82 *S*. *agalactiae* genomes analysed by Piggy (57).

Frequency analysis of IGRs in *S*. *agalactiae* pan-genome showed that the number of IGRs shared across all strains was smaller than core protein-coding genes, whereas strain-specific IGRs was much higher than protein-coding strain-specific genes (Fig. 2C). IGR analysis with Piggy excludes IGRs that are less than 30 bp which may result in fewer IGRs than protein-coding genes in core regions (57). Most IGRs identified in the pan-genome belonged to either core genes or strain-specific genes (Fig. 2C) in line with previous findings in *Staphylococcus aureus* ST22 and *Escherichia coli* ST131 where similar distributions of IGRs were detected (57). The gradients of the accumulation curves for both protein-coding genes and IGRs are still strongly positive therefore the *S*. *agalactiae* pan-genome remains open.

### 3.3. Aquatic serotype Ib have a reduced repertoire of virulence factors

Almost all genes classed as adhesins, involved in immune evasion and host invasion, and most of the toxin-related genes found in terrestrial isolates were absent from serotype Ib aquatic isolates (Fig. 3). Rosinski-Chupin et al. (2013) reported ~60% of the virulence genes found in human strains were present in a serotype Ib GBS strain from fish. We found that CAMP factor gene *cfa/cfb* was present in all strains including serotype Ib ST-260, 261 and 552 isolates, but CAMP reaction was previously reported to be negative for ST-260 and 261 strains (13, 28). These authors identified that CAMP factor gene in ST-261 is disrupted while the gene in ST-260 is unaltered but the level of gene expression may be too low to detected by the test (13). Most of the genes in *cyl* locus have been lost from Ib piscine strains and only *cylB*, an ABC ATP binding cassette transporter was present in ST-260 and 552 (Fig. 3). The *cyl* locus is responsible for hemolytic activity and production of pigment via co-transcription of *cylF* and *cylL* (73). In ST-261, the *cyl* gene cluster is replaced by a genomic island resulting in loss of hemolytic activity (13). Of particular relevance to virulence and antigenic diversity, transmembrane immunoglobulin A-binding C protein beta-antigen *cba* was absent in all aquatic isolates (Fig 3). This gene has been reported in type Ib and Ia GBS previously (74), is implicated in virulence in neonates and is up regulated in GBS serotype Ia strain A909 in response to human serum (74–76). In contrast, capsule-related genes were largely conserved and associated with serotype (Fig. 3) supportive of the major role of capsule in virulence of fish pathogenic streptococci (77, 78).

**Figure 3.**
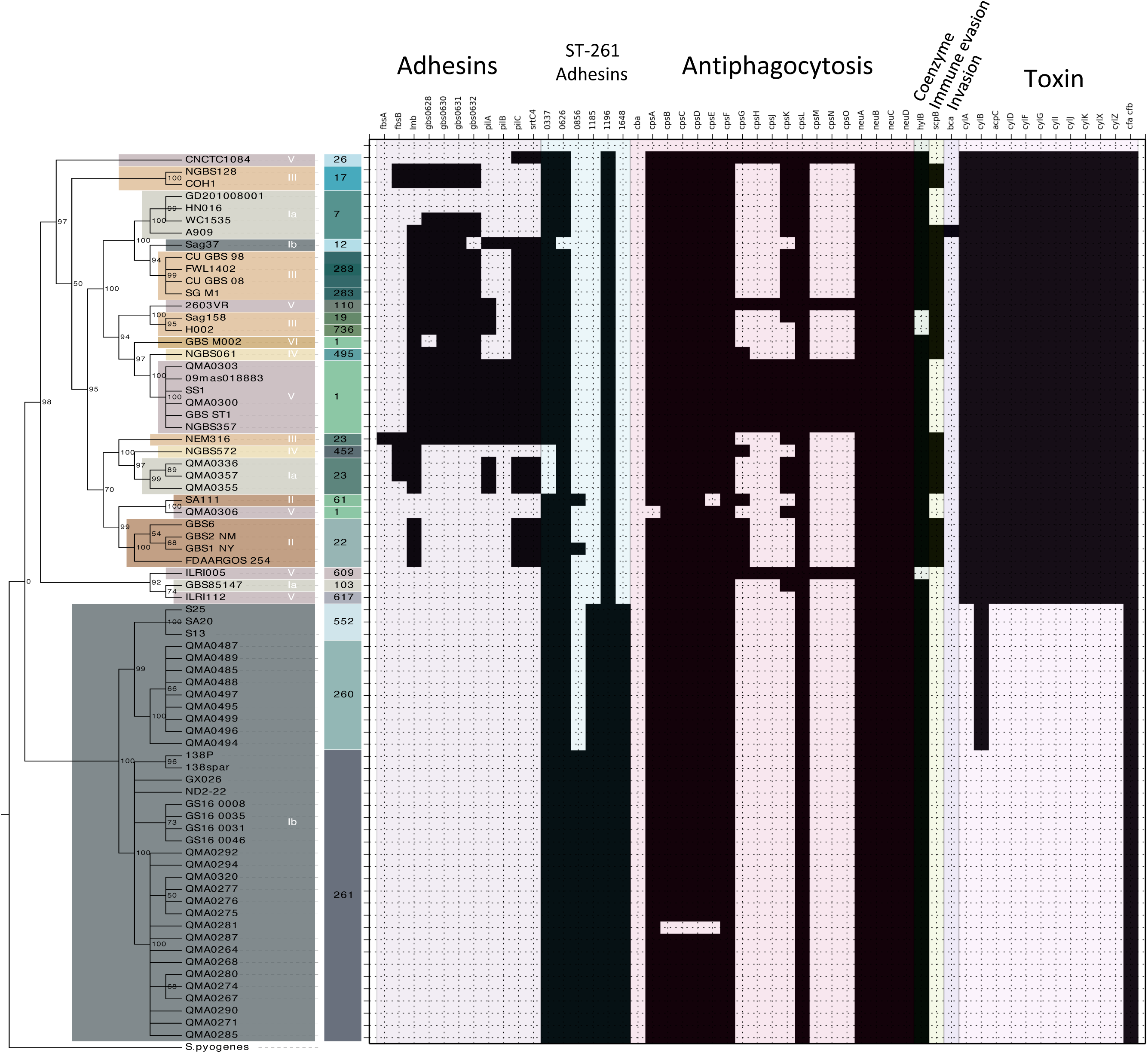
Virulence gene presence and absence in *S*. *agalactiae*. Genes were identified from VFDB to create a *S*. *agalactiae* database for comparison of 82 strains by BLAST using SeqFindr with default settings (95% identity cut off).

Although serotype Ib aquatic isolates have lost the majority of virulence factors found in terrestrial isolates, most contained six sequences that were identified as probable adhesins by homology (Fig. 3). These ST-261 adhesins (named 0337, 0626, 0856, 1185, 1196 and 1648 based on position in the annotated ST-261 genome from QMA0271) were fully conserved amongst aquatic serotype Ib ST-261 (Fig. 3). Moreover, ST-261 adhesin 0856 was only present in ST-261 and two serotype II isolates, GBS1-NY and SA111 strains (Fig. 3). ST-261 adhesins 1185 and 1648 were shown to be unique to aquatic serotype Ib isolates, being absent from Ia fish isolates and all terrestrial strains (Fig. 3). The analysis indicated that ST-261 adhesins 0337, 0626 and 1196 were well conserved across most of the isolates regardless of origin but 1196 was the only adhesin-like gene present in all strains analysed (Fig. 3). ST-261 adhesin 0626 was also ubiquitous but contained deletions in Sag37 isolate (Fig. 3). Some terrestrial strains, such as QMA0355, 0357, 0336 and NGBS572 lacked adhesin 0337 (Fig. 3).

As capsular polysaccharide is a requirement for full virulence in several fish pathogenic streptococci (77, 78) and the major antigen in GBS (79–81), further analysis of the serotype Ib *cps* operon was conducted. Within the serotype Ib lineage, the capsular polysaccharide operon was well conserved (Fig. 4). However, QMA0281 from grouper had a deletion at position 341 in *cpsB*, encompassing *cpsC*, *cpsD* and *cpsE* (Fig 4), resulting in a chimeric ORF. Moreover, an insertion in a TA repeat at position 557 in *cpsH* resulted in an early stop codon marginally reducing gene size (Fig. 4). QMA0368 had a TT insertion at nucleotide position 745 in *cpsE* resulting in insertion of an early stop codon and truncation of the gene (Fig. 4). Deletion of this region that includes the priming glycosyl transferase for capsular biosynthesis results in loss of capsule, attenuated virulence and modified pathology in the fish pathogen *S*. *iniae* (82). Buoyant density analysis in Percoll indicated that QMA0281 is also deficient in capsule (not shown). *cps* operon SNPs amongst the Australian serotype Ib isolates were rare (Fig. 4). Indeed only 1 SNP *neuA* separated QMA0271 from the 1988 tilapia isolate from Israel suggesting very little evolution of the *cps* operon since introduction of the lineage (Fig. 4). This may reflect a well-adapted capsule for colonization naïve hosts as the Australian isolates were from wild fish and captive stingrays recently after capture and transport thus placing little selective pressure for novel capsular sequence types. Immunity in fish drives evolution of novel capsular sequence types in *S*. *iniae* but these reported cases were all in high intensity aquaculture with occasional use of autogenous vaccination and opportunity for development of cps-specific immunity (82). This may explain the relatively high number of SNPs in genes encoding sugar and sialic acid modifying enzymes amongst the isolates from tilapia farmed in Honduras relative to the other isolates examined, including those from tilapia, as autogenous vaccination is occasionally used on farms where isolates were sourced, and may apply selective pressure favouring modified polysaccharide capsule.

**Figure 4.**
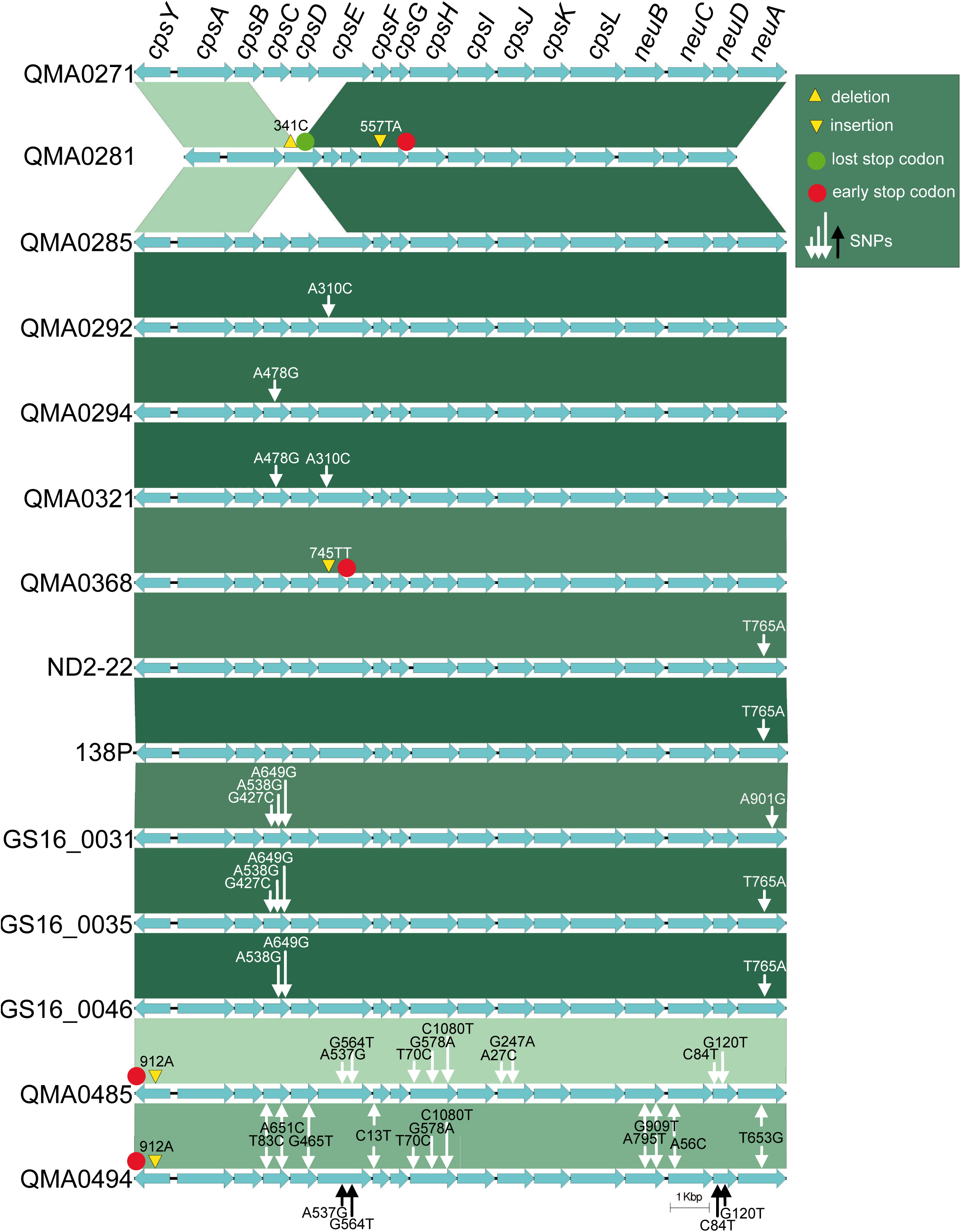
The capsular polysaccharide (cps) operon of the ST-261 lineage. The operon was identified in Genbank files manually and then compared by BLAST using EasyFig (89)

## Conclusions

The clade of aquatic *S*. *agalactiae* serotype Ib including ST-260, ST-261 and ST-552 is a highly adapted fish pathogen with a substantially reduced genome compared to all other serotypes from terrestrial mammalian, reptile and fish hosts. These variants were originally identified as *S*. *difficile* due to their impoverished growth on laboratory media and hence difficulty in isolating from diseased fish (70). The species was reassigned to serotype Ib GBS a few years later (69) but the recent discovery of the major reduction in genomes size of ST-261 serotype Ib (13) explains the marked phenotypic difference that merited early phenotypic assignation to its own species. Other serotypes have been isolated from fish, notably serotype Ia and serotype III, but these seem to arise from terrestrial transfer rather than being retained amongst the aquatic host population (28). In contrast, serotype Ib has only been isolated from fish and stingrays, appears to be well-adapted and is likely to have been transferred internationally via trade in domesticated tilapia, evidenced here by the very close relationship (only a handful of SNPs) between a strain isolated from tilapia in 1988 in Israel and those found in fish and stingrays in Australia since 2008, and in tilapia in the USA, China and Ghana. The ST-261 lineage in Australia likely arrived with several introductions of tilapia in the 1970s and 1980s. Tilapia are classed as a noxious pest in Australia and import was banned, but not before several lines became established throughout tropical and subtropical freshwater habitats in Queensland (71). Although ST-261 serotype Ib GBS has not been isolated from farm fish in Queensland in spite of proximity of freshwater farms to tilapia-infested creeks, this clade of GBS is a substantial problem in farmed tilapia globally. The cohort of putative adhesins identified here to be conserved through all fish pathogenic serotypes (including Ia and III in addition to Ib) may be promising candidates for cost-effective cross-serotype protective vaccines for aquaculture and are worthy of future research.

## Acknowledgements

This work was funded by the Fisheries Research and Development Corporation Aquatic Animal Health Subprogram Project 2010/34 awarded to ROB and ACB, and the Australian Research Council Linkage Program LP130100242 awarded to ACB, MJW and SB.

